# Single-cell RNA-seq of rheumatoid arthritis synovial tissue using low cost microfluidic instrumentation

**DOI:** 10.1101/140848

**Authors:** William Stephenson, Laura T. Donlin, Andrew Butler, Cristina Rozo, Ali Rashidfarrokhi, Susan M. Goodman, Lionel B. Ivashkiv, Vivian P. Bykerk, DE Orange, Robert B. Darnell, Harold P. Swerdlow, Rahul Satija

**Author notes:** these authors contributed equally (William Stephenson, Laura T. Donlin, Andrew Butler, Harold P. Swerdlow, Rahul Satija).

## Abstract

Droplet-based single cell RNA-seq has emerged as a powerful technique for massively parallel cellular profiling. While these approaches offer the exciting promise to deconvolute cellular heterogeneity in diseased tissues, the lack of cost-effective, reliable, and user-friendly instrumentation has hindered widespread adoption of droplet microfluidic techniques. To address this, we have developed a microfluidic control instrument that can be easily assembled from 3D printed parts and commercially available components costing approximately $540. We adapted this instrument for massively parallel scRNA-seq and deployed it in a clinical environment to perform single cell transcriptome profiling of disaggregated synovial tissue from a rheumatoid arthritis patient. We sequenced 8,716 single cells from a synovectomy, revealing 16 transcriptomically distinct clusters. These encompass a comprehensive and unbiased characterization of the autoimmune infiltrate, including inflammatory T and NK subsets that contribute to disease biology. Additionally, we identified fibroblast subpopulations that are demarcated via *THY1* (CD90) and *CD55* expression. Further experiments confirm that these represent synovial fibroblasts residing within the synovial intimal lining and subintimal lining, respectively, each under the influence of differing microenvironments. We envision that this instrument will have broad utility in basic and clinical settings, enabling low-cost and routine application of microfluidic techniques, and in particular single-cell transcriptome profiling.

## Introduction

The complex architecture and associated higher-order function of human tissues relies on functionally and molecularly diverse cell populations. Disease states represent significant perturbations to cellular heterogeneity, with tissue-resident cells acquiring altered phenotypes and circulating cells infiltrating into the tissue. Therefore, defining the cellular subsets found in pathologic tissues provides insights into disease etiology and treatment options. Traditional methods such as flow cytometry, which require *a priori* knowledge of cell type-specific markers, have begun to define this landscape, but fall short in comprehensively identifying cellular states in a tissue, with particular difficulty detecting novel or extremely rare subpopulations.

Technological advancements in automation, microfluidics, and molecular barcoding schemes have permitted the sequencing of single cells with unprecedented throughput and resolution^1–4^. In particular, recent studies featuring analysis of 10^4^−10^5^ single cells have enabled unbiased profiling of cellular heterogeneity, where entire tissues can be profiled without advance enrichment of individual cell types^1,5,6^. In spite of this progress, technological advances can be slow to permeate into resource-limited clinical arenas due to a variety of reasons related to cost, personnel requirements, space or infrastructure. Specifically, a major barrier to widespread adoption of droplet microfluidic techniques is the lack of cost-effective and reliable instrumentation^7,8^. Microfluidic experiments are typically performed using commercial instruments which are expensive and often configured for a single purpose, or custom research instrument setups which are comprised of multiple pieces of equipment and rarely portable. Particularly in clinical settings, microfluidic instrumentation is not always proximal to the site of cell sample generation requiring transport to external sites or cell preservation, both of which can alter cellular transcriptomes or result in extensive cell death^6,9^.

To address these short-comings and provide a low-cost option for single-cell transcriptome profiling, we have developed a portable instrument for performing single-cell droplet microfluidic experiments in research and clinical settings. Recent microwell-based transcriptome profiling approaches have been shown to be advantageous for low-cost portable transcriptome profiling^10–12^, however some of these techniques are challenging to perform and or require extensive chemical modification to fabricate the devices. Additionally, the fixed architecture of microwell (partitioning) microfluidic devices dictates their use for specific applications. In contrast, the platform presented here is easy to use and can be implemented for a variety of droplet microfluidic (partitioning) or continuous phase microfluidic based experiments. Potential applications of this system include recent work profiling immune repertoires from hundreds of thousands of single cells^13^ and combined single-cell transcriptome and epitope profiling^14^ in addition to ddPCR^15^, ddMDA^16^, hydrogel microsphere fabrication for 3D cell culture^17,18^, chemical microfluidic gradient generation^19^ and microparticle size sorting^20–22^. The instrument is comprised of electronic and pneumatic components affixed to a 3D printed frame. The entire system is operated through software control using a graphical user interface on a touchscreen. Requiring only a standard wall power outlet, the instrument has an extremely small footprint; small enough to fit on a bench top or in a biocontainment hood. The total cost of materials to construct an instrument is approximately $540. (Supplementary Table 1) This represents an approximately 20-fold reduction in cost for a research-level, syringe-pump based microfluidic setup, and a 200-fold reduction in cost for a commercial microfluidic platform.

We applied the microfluidic control instrument in conjunction with the Drop-seq technique^1^ to perform unbiased identification of transcriptomic states in diseased synovial tissue, which becomes highly inflamed in rheumatoid arthritis (RA) and drives joint dysfunction. RA is a common autoimmune disease affecting approximately 1% of the population. While the cause of RA is not precisely known, disease etiology is hypothesized to originate from a combination of environmental and genetic factors^23,24^. RA affects the lining of the joint; the synovial membrane, leading to painful inflammation, hyperplasia, and joint destruction. RA is clinically characterized by multiple tender and swollen joints, autoantibody production (rheumatoid factor and anti-citrullinated protein antibody or ACPA) in addition to cartilage and bone erosion^25^. Unlike other tissue membranes with an epithelial layer, the synovial lining is composed of contiguously aligned fibroblasts and macrophages 2–3 cells deep^26^. In RA, the membrane lining is expanded to 10 – 20 cells deep and synovial fibroblasts assume an aggressive phenotype marked by the expression of disease relevant cytokines, chemokines and extracellular matrix remodeling factors.^27–29^ The sublining is marked by an accumulation of lymphocytes, macrophages, and dendritic cells amidst the subintimal synovial fibroblasts. Pioneering studies have uncovered heterogeneity in fibroblast morphology^30^ and phenotype^31,32^, observing differences in activation state and invasive behavior^33,34^. In addition, in situ hybridization has identified non-uniform activation of inflammatory drivers and matrix metalloproteinases^35,36^, motivating the use of our unbiased approach to catalogue fibroblast subpopulations, and molecular markers which define them.

Here we describe the design of a novel microfluidic control instrument that can be assembled with 3D printed and commercial components at low cost, is fully portable, and functions as a reliable and flexible droplet generator. We adapted this device to perform massively parallel single cell RNA-seq (Drop-seq), observing metrics and performance that were indistinguishable from a research level Drop-seq setup. We deployed this instrument to a hospital laboratory to profile 8,716 single cells from the synovial tissue of an RA patient. To our knowledge, this represents the first unbiased ‘atlas’ of hematopoietic and fibroblast transcriptional subtypes from scRNA-seq of autoimmune disease tissue. We identified 16 subpopulations, including both abundant and rare groups that contribute to disease biology. We also define novel cellular subsets of synovial fibroblasts, which were validated and used to leverage additional insights by immunofluorescence and flow cytometry. The deconvolution of cellular complexity in a diseased tissue by this portable device provides a template for the application of droplet-based single-cell transcriptome profiling for routine clinical analysis.

## Results

### Development of a portable, low-cost droplet microfluidic control instrument

To perform single-cell transcriptome profiling experiments in clinical settings at low-cost, the components of a standard Drop-seq setup were replaced with alternative miniature components and packaged onto a multitiered 3D printed frame. (Figure 1a,b,c, Supplementary Figure 1) For example, syringe pumps in a standard Drop-seq setup (which provide a means for fluid flow through a microfluidic chip) were replaced with components such as a micro air-pump, regulators, and micro solenoid valves. These components are just as effective for providing adequate fluid flow, in a much smaller footprint and at significantly lower cost. Stirring of barcoded microparticles is achieved through actuation of a stepper motor affixed with a permanent magnet at the end of 3D printed shaft. Rotation of the motor shaft locally inverts a magnetic field thereby tumbling a magnetic stir disc in the microparticle fluid reservoir. A custom printed circuit board (PCB) was designed to interface the electronic and pneumatic components of the instrument to a single board computer (Raspberry Pi). Further critical components of the instrument include pressure sensors for optimal flow rate determination, micro solenoid valves for on-demand pressure actuation, and a microscope for real-time experiment monitoring. The microscope is comprised of an inexpensive 5-megapixel CMOS camera coupled with a laser diode collimating lens. This provides sufficient magnification operating in fixed-focus mode to view the microfluidic channels with the ability to resolve single cells. (Figure 1d, Supplementary Video 1) The instrument is operated through a custom graphical user interface on a touchscreen. All components were affixed to a 3D printed frame measuring approximately 21 cm by 20 cm and 9 cm tall. (Figure 1b) For Drop-seq experiments, fluorinated oil, cells, and barcoded microparticles are pipetted into fluid reservoirs situated at the rear of the instrument. Custom pressure caps seal the vial and tubing connections are made to a microfluidic chip situated on the top of the instrument above the microscope camera. The small footprint of the device permitted use in clinical laboratory space requiring only a standard wall outlet for power.

**Figure 1.**
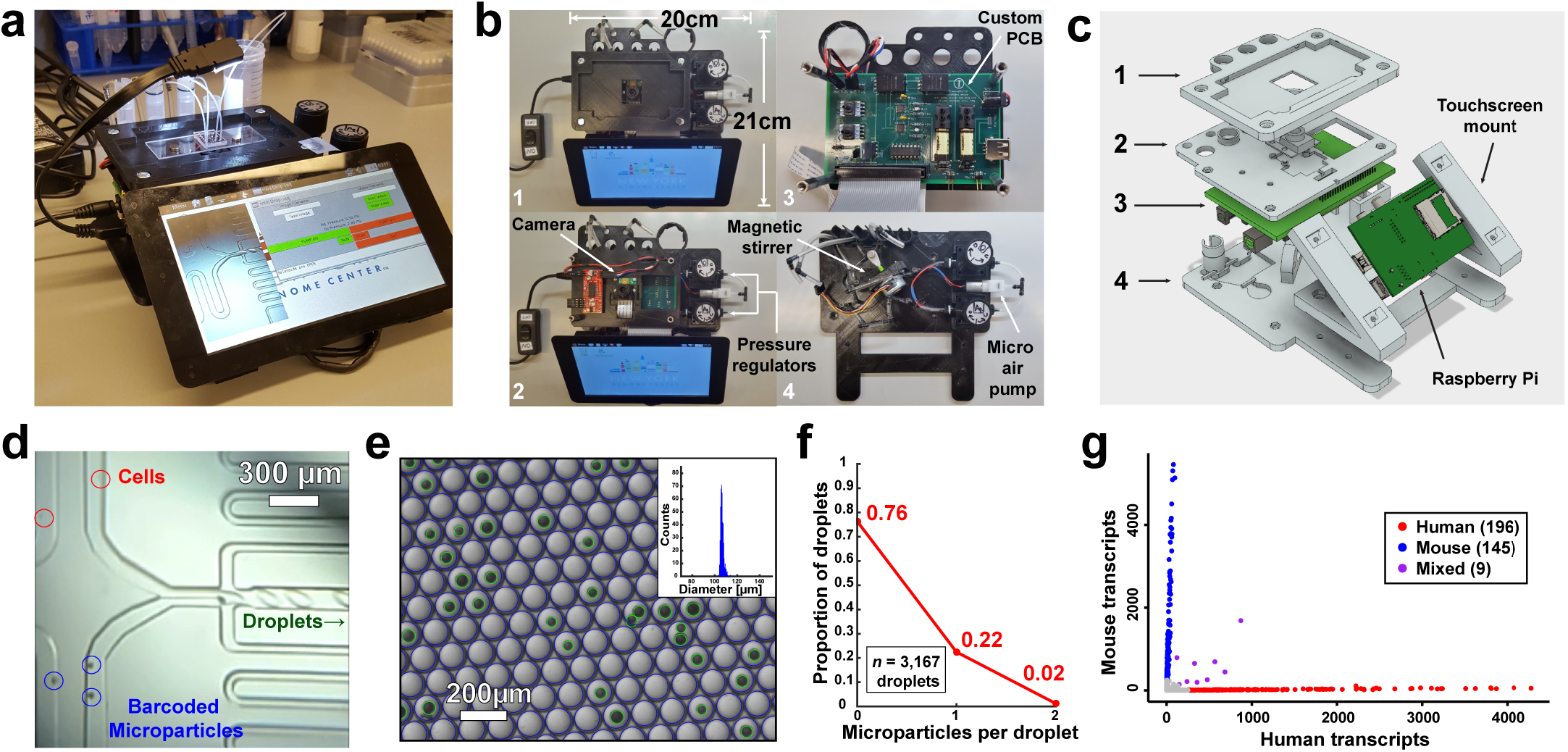
Microfluidic control instrument design and validation. a) Picture of the microfluidic control instrument performing a Drop-seq run. b) Top down views of multi-tiered instrument. Levels 1–4 reveal assorted components for instrument operation. c) 3D rendering of the instrument with levels corresponding to those in b). Components in light gray are 3D printed. d) Microscope image screen capture directly from the instrument. Cells and barcoded microparticles are visualized easily on the screen. e) Microscope image of droplets output from the instrument. Droplets and microparticles are detected via image analysis software as blue circles and green circles respectively. Inset: droplet diameter distribution histogram. f) Microparticle loading distribution into droplets as measured via automated image analysis is consistent with Poisson loading. g) Species mixing experiment using mouse (3T3) and human (HEK293) cells.

### Instrument validation and operation

To validate the design and operation of the instrument we first assessed the droplets produced by the device in conjunction with a slightly modified Drop-seq microfluidic chip. (Supplementary Figure 1d) Droplets produced using the instrument displayed a high degree of uniformity (diameter = 105 +/− 3 μm) across multiple microfluidic chips and instruments at identical operating pressures indicating stable and reproducible flow patterns of the assembly. (Figure 1e) Next, the loading of microparticles (tested at an optimized concentration) into droplets was assessed using a custom written MATLAB image analysis script to measure the number of empty, singly occupied, and doubly occupied droplets. The resulting microparticle loading profile followed a Poisson statistical distribution, which is expected for a stochastic loading process such as encountered here. (Figure 1f) Finally, to validate single-cell encapsulation, we performed a species-mixing experiment in which approximately equal numbers of HEK293 (human) cells and NIH 3T3 (mouse) cells were combined in a single run, followed by shallow sequencing (average of 2,017 reads/cell, 1,187 unique molecules (UMI)/cell) (Figure 1g). We observed low doublet rates (2.57%), and high species-specificity across singlets (98.2%), consistent with technical metrics for Drop-seq.

### Transcriptomic profiling of synovial tissue

After validation of the instrument, we turned to profiling the inflammatory cellular milieu in synovial tissue extracted from the knee of a seropositive RA patient (Figure 2a). Histologically the tissue displayed characteristics of extensive inflammation, including synovial lining hyperplasia (black arrow) and dense leukocyte infiltrations in the sub-lining (blue arrow) (Figure 2b). Fibroblast morphology within the sub-lining varied widely in the intervening space suggesting that subpopulations of fibroblasts may exist in heterogeneous micro-niches. Immediately after surgery, a portion of the recovered joint tissue was processed using an optimized disaggregation protocol to generate a single cell suspension. Cells were counted, resuspended for optimal single cell loading into droplets, and immediately pipetted into the appropriate fluid reservoir of the instrument to run through the Drop-seq protocol. Briefly, following encapsulation in droplets, cells are lysed and mRNAs hybridized to microparticles undergo reverse transcription in bulk to generate stable cell-barcoded cDNAs as previously described^1^. The total time starting with sample extraction from the patient to initiation of the microfluidic instrument is approximately 1.5 hours, obviating the need for cell preservation. The instrument processes 1 mL of cells at a concentration of 150 – 200 cells/μl in about 35 minutes, generating over one million droplets at a generation rate of approximately 660 Hz. We performed two replicate runs on identical instruments simultaneously to obtain a total of 8,716 single cell transcriptomes, sequenced to an average read depth of 30,443 reads/cell, and detecting an average of 1,903 unique molecules per cell. This corresponds to a profiling throughput (per individual instrument) of approximately 7,500 cells/hour. Therefore the throughput, cell capture, and input requirements are in line with a research Drop-seq setup^1^.

**Figure 2.**
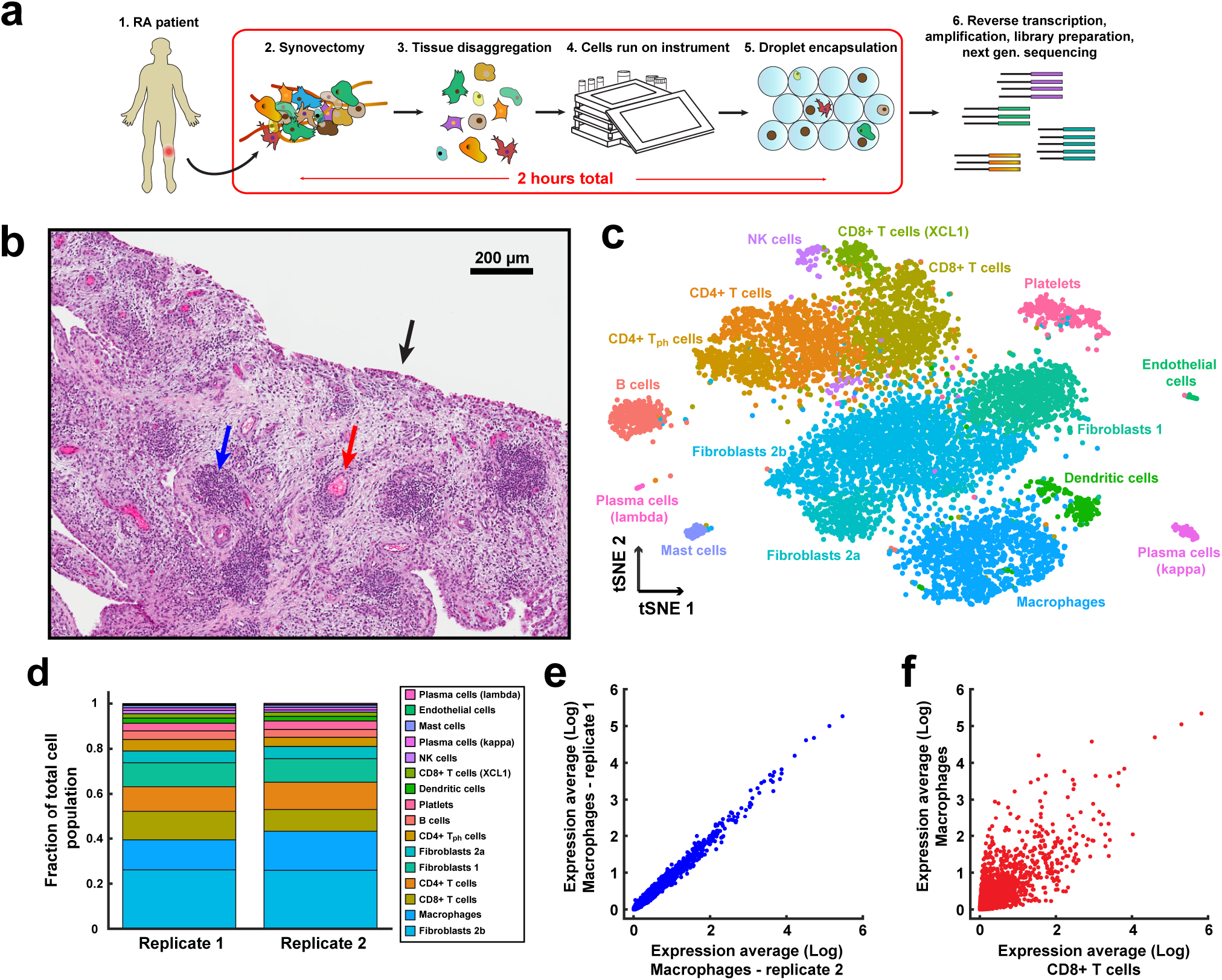
RA sample workflow and histology, single-cell unsupervised clustering and analysis. a) Sample workflow from operating room to sequencing. Preparation of single cells into droplets with barcoded microparticles is performed in about 2 hours. b) H&E stain of synovial tissue from the patient. The synovial lining is indicated by the black arrow. An example of vasculature is indicated by a red arrow. The blue arrow denotes a dense lymphocyte infiltrate. c) Unsupervised graph-based clustering of single-cell RNA-seq, visualized using t-distributed stochastic neighbor embedding (tSNE). Each point represents a single cell (droplet barcode). d) Fraction of total cells present in each cluster, subdivided by replicate. e) Bulk expression (‘in silico average’) comparisons across macrophages from each replicate. f) Expression comparison across combined CD8+ T cell and macrophage population from both replicates.

We applied our previously developed graph-based clustering procedure^10,37^, to partition cells into 16 distinct subpopulations, which we visualized using t-distributed stochastic neighbor embedding (t-SNE) (Figure 2c). While the clustering was unsupervised, differential expression revealed combinations of known markers that could be used to confidently assign subpopulations to broad categories. For example, we observed 13 immune populations that broadly expressed *PTPRC* (CD45) and three fibroblast populations, expressing uniform high levels of *COL1A2*. Similarly, as we explored further within immune cells, we identified clear markers of known subtypes types, including canonical macrophage markers *(MARCO)*, T cell *(CD3)* and B cell (*MS4A1*, CD20) markers (Figure 3).

**Figure 3.**
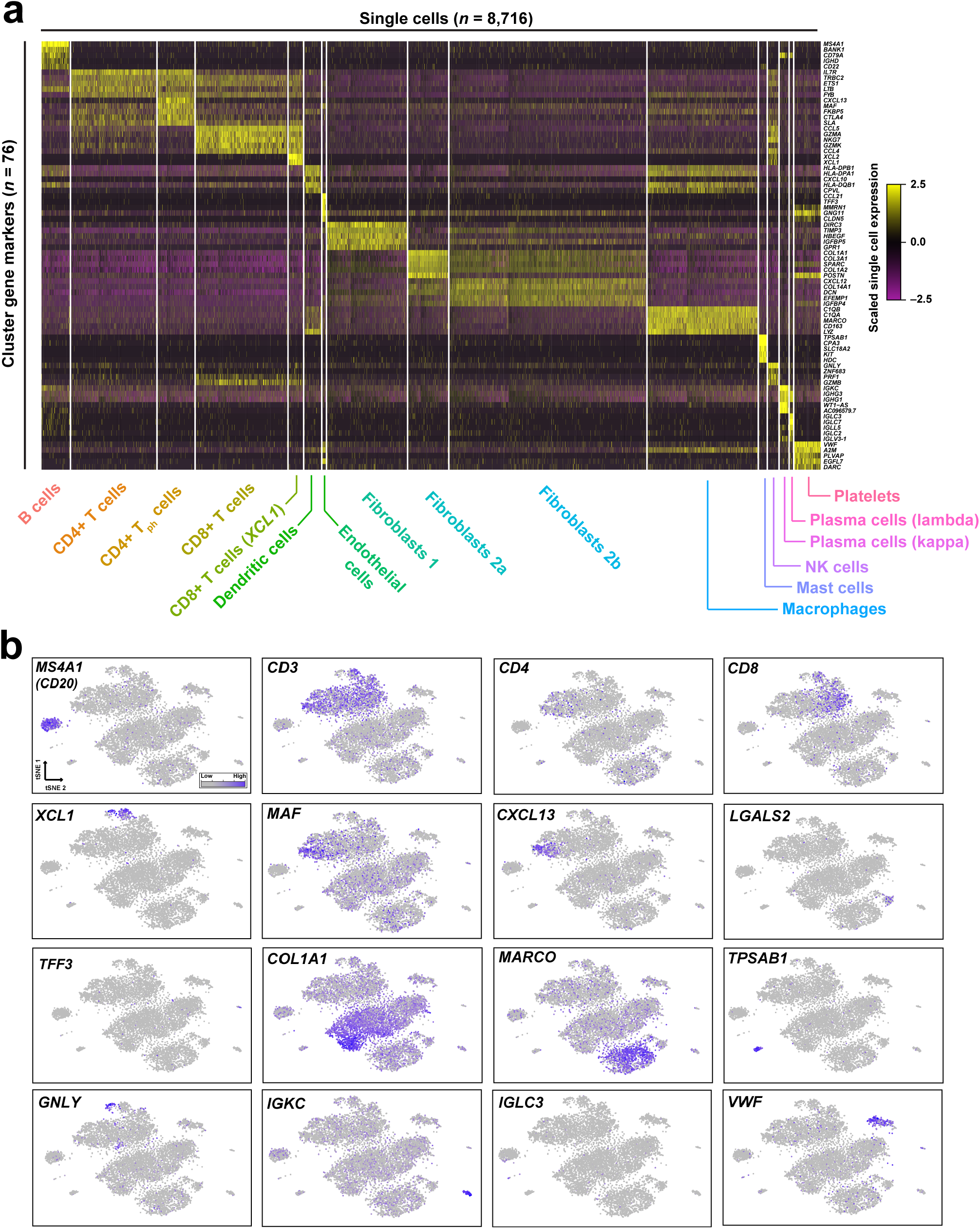
Transcriptomic markers of gene expression for individual clusters. a) Single cell expression heatmap displaying up to five transcriptomic markers for each cluster, based on differential expression testing. b) Gene expression for canonical marker genes, overlaid on the tSNE visualization. A list of transcriptomic markers for each cluster is provided in Supplementary Data 1.

The proportion of cells mapping to each of the 16 clusters was tightly conserved between replicate experiments (R=0.98). Additionally, we compared averaged expression levels for cells in the same cluster across replicates. For example, global macrophage transcriptomes were highly reproducible between replicates (R=0.99) (Figure 2e), but transcriptomes for different cell populations were widely divergent as expected (macrophage/CD8+ T cell R=0.82) (Figure 2f). These results demonstrate the reproducibility of the overall workflow. Additionally, our reproducible and quantitative ‘in silico’ bulk transcriptomes offer an alternative to traditional bulk RNA-seq on sorted populations, as our procedure requires no sorting, and can derive averages for all 16 populations simultaneously.

To our knowledge, this single-cell dataset represents the first unbiased and comprehensive ‘atlas’ of cellular subpopulations present in human autoimmune disease tissue. Below, we summarize both abundant and rare cell states in our data, with unbiased markers shown in Figure 3 and Supplementary Data 1. We highlight particular subtypes of lymphocyte and myeloid cells that have not been previously identified in healthy PBMCs, as well as unexpected transcriptomic heterogeneity within fibroblast populations.

## Unsupervised taxonomy of cellular states in synovial tissue

We identified 8 lymphocyte subpopulations corresponding to heterogeneous groups of T, B, and NK cells. T cells (CD3+) were grouped into CD4+ (helper) and CD8+ (cytotoxic) subpopulations based on canonical markers. Within the CD4+ T helper cell population we detected a distinct subset marked by high levels of *MAF, CXCL13 and PDCD1* (PDL1), which has not been previously identified in previous single cell RNA-seq studies of human PBMCs^4,10^. However, a recent CyTOF analysis of RA synovial tissue identified a population with consistent markers, representing an RA synovial “peripheral T helper cell” (T_PH_) that may support B cell activity and antibody production in this non-lymphoid tissue^38^ (Figure 2c). Pathway enrichment analysis tailored to single cell data^39^ identified functional modules up-regulated specifically in these cells, including the regulation of B cell proliferation and T cell chemotaxis (Supplementary Figure 2a), supporting these functional analyses, and demonstrated our ability to identify cellular phenotypes that are unique to diseased tissue.

We also observed further cellular heterogeneity within the CD8+ and NK lymphocyte subsets. In particular, for both classes we observed subsets of cells expressing extremely high levels of the cytokines XCL1 (lymphotactin) and XCL2, which have previously been demonstrated to be present at higher levels in synovial RA tissue. While CD8+ T cells are known to express XCL1^40^, our observation that this is restricted to only a specific subpopulation (Figure 3a,b) may suggest a functionally important role for this group, particularly in stimulating the trans-migration of primed lymphocytic subsets, or modulating matrix metalloproteinase expression in synovial fibroblasts^41^. For the NK cells (uniformly expressing GNLY), principal component analysis revealed a subpopulation expressing similarly high levels of XCL1 and XCL2, while also down-regulating cytotoxic genes (*PRF1*) and *FCGR3A* (CD16), representing a bifurcation between CD16^+^CD56^bright^ and CD16^−^CD56^dim^ subsets (Supplementary Figure 2b). Notably, while CD56^bright^ cells are rare in healthy tissue and have not been identified in scRNA-seq analyses of PBMCs, we detect them at 37% frequency here, consistent with previous reports that the presence of this subset is enriched within RA tissue^42^.

We also characterized B cell populations (*MS4A1*+) (also known as CD20), as well as terminally differentiated populations that secrete high levels of immunoglobulins (*IGHG4*+). Our single-cell dataset indeed distinguished two distinct populations of plasma cells, again not previously observed in existing scRNA-seq data of PBMCs, based on antibody light chain usage (IgA kappa+ vs. IgA lambda+). This enabled us to calculate a kappa/lambda ratio based on single cell proportions, that was conserved across replicates (2.5 replicate 1, 2.8 replicate 2). (Figure 3b). Finally, we also identified four non-lymphocytic hematopoietic subpopulations, including mast cells (*TPSAB1*+), macrophages (*MARCO*+), dendritic cells (*HLA-DRB5*+), and platelets (*VWF*+). Taken together, these clusters represent an unbiased and detailed characterization of tissue-resident immune cells from inflamed synovial tissue, including both abundant and rare populations that contribute to disease biology.

## Identification and validation of heterogeneous fibroblast subtypes

Non-hematopoietic cells were composed primarily of fibroblasts based on a commonly expressed set of genes consistent with the fibroblast lineage such as *COL1A2, COL3A1* and *CLU* (Figure 3a). While immune cell subsets can be defined based on canonical marker expression, potential source of cellular heterogeneity in fibroblasts are poorly understood, despite their strong implication in inflammatory diseases^26–28^. Our unbiased clustering returned three fibroblast subpopulations (Figure 3a, 4a-b). These represented two groups of fibroblasts with distinct bifurcations in marker expression (Fibroblast 1 vs. Fibroblast 2), as well as a further a subdivision of the latter (Fibroblast 2a vs. Fibroblast 2b) representing more quantitative differences in gene expression. Genes differentially expressed between the subsets included known drivers of RA biology, including cytokines (CXCL12), matrix metalloproteinases (MMP2, MMP3), in addition to a subset of surface protein markers (i.e. CD55; CD90) (Fig. 4c).

**Figure 4.**
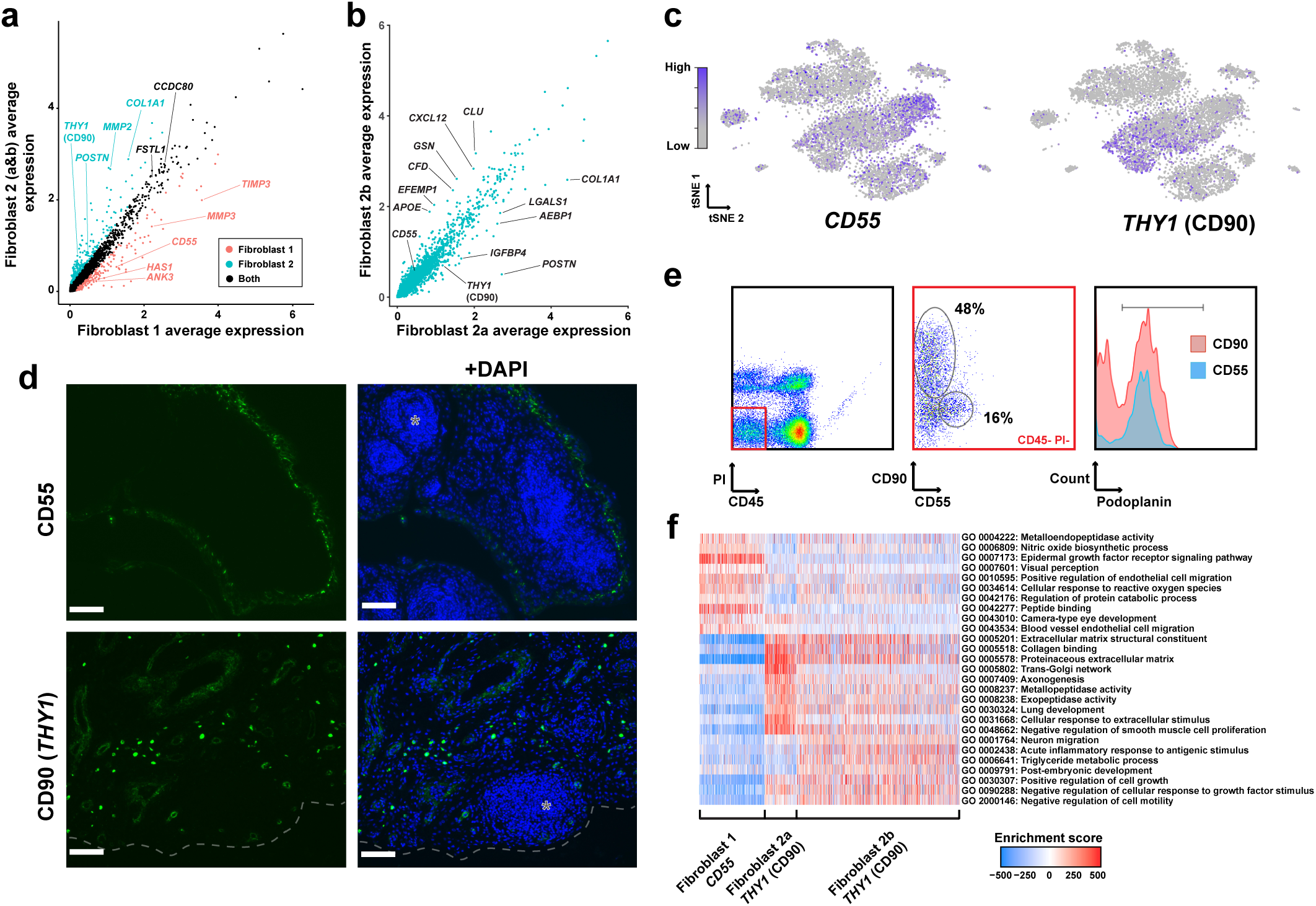
Identification of synovial fibroblast subtypes. a) Bulk expression (‘in silico average’) comparison of fibroblast populations (1 vs. 2a&b). Genes up-regulated in fibroblast population 1, based on differential expression analysis with Bonferroni-corrected p<0.05, are indicated in red and genes expressed predominantly in fibroblast population 2 are indicated in teal. b) Expression comparison across fibroblast sub-populations 2a and 2b. c) *CD55* and *THY1* (CD90) expression across the global tSNE. d) CD90 and CD55 localization in RA synovial tissue. Paraffin-embedded synovial tissue was sectioned and assayed for target markers by immunofluorescent staining with antibodies for CD90 (green) or CD55 (green) and counterstained with DAPI (blue). Lymphocyte infiltrates are denoted by grey asterisks. Images were acquired at 20x magnification. Scale bar is 100 μm. e) Cell surface expression of CD90, CD55 and podoplanin in synovial tissue. Synovial cell suspensions were assayed for target markers by flow cytometry and stained with CD45, CD55, CD90, podoplanin and PI. Synovial cells were gated on the CD45− PI-population and analyzed for the proportion of CD90+ and CD55+ cells. Relative podoplanin cell surface expression was analyzed in CD90+ (red) and CD55+ (blue) as shown in histograms. f) Pathway and gene set overdispersion analysis on the three fibroblast populations identified from unbiased clustering of single cell RNA-seq data. Enrichment score corresponds to each cells’ first principle component loading from pathway analysis as computed in *pagoda*.

We next looked to validate the major separation of fibroblast subsets, and to test the specificity of the putative markers using complementary techniques. To this end, paraffin-embedded tissue blocks for this tissue were sectioned and analyzed by immunofluorescence with antibodies against subset-specific markers (Figure 4d). Importantly as this approach examines the cells and markers within the intact tissue, it eliminates dissociation-induced artifacts and potentially informs on anatomic localization within the tissue. Interestingly CD55 (Fibroblast 1 marker) predominantly stained in the synovial lining (Figure 4d). Distinctly, CD90 antibodies (Fibroblast 2 marker) labeled cells in the sublining regions, with intense staining around presumed small vessels and intermediate staining that encircled wider rings around larger vessels (Figure 4d).

Furthermore, a flow cytometric analysis of non-hematopoietic viable cells from this tissue demonstrated that CD90 and CD55 antibodies stained independent cell populations (Figure 4e, middle panel). The CD55+ cells were also largely positive for the common fibroblast marker podoplanin, while the CD90+ non-hematopoietic cells separated into a podoplanin-positive (fibroblasts) and -negative population (Fig. 4e, right panel). The CD90+ CD45− PDPN− population likely represents endothelial cells^43^. These data further indicate the unique nature of fibroblast subsets in RA synovial tissue and suggest an approximate ratio of 3:1 sublining CD90+ versus lining CD55+ fibroblasts, consistent with our scRNA-seq data.

The distinct anatomical distribution of Fibroblast 1 and 2 populations hereafter referred to as CD55+ lining and CD90+ sublining fibroblasts, respectively, implicate putative functional differences. CD55+ fibroblasts locate to the intimal lining, which is responsible for the generation and turnover of synovial fluid. Importantly, hyaluronan synthase 1 (*HAS1*) expression was enriched in CD55+ lining fibroblasts. (Figure 4a) As hyaluronan represents the most abundant macromolecule in synovial fluid, this suggests these cells function within the lining to produce synovial fluid components. Pathway and gene-set enrichment analysis revealed hierarchical relationships between fibroblast subpopulations that were consistent with transcriptomic similarities, and also suggested heterogeneous pathway activation between these groups. For example, CD55+ fibroblasts exhibited stronger activation of epidermal growth factor signaling and metalloendopeptidase activity, while both CD90+ fibroblast groups were enriched for modules associated with the production and interaction with the extracellular matrix. Between the CD90+ groups, we observed more quantitative differences in the activation of acute inflammatory responses, supporting the role of these sublining fibroblasts interacting with lymphocytic infiltrates. Collectively, distinct anatomic locations, cell surface staining and transcriptomic differences confirm the independent nature of these synovial fibroblast subsets.

## Discussion

In this study, we developed a low-cost, portable microfluidic control instrument to perform droplet-based singlecell transcriptomic profiling in a clinical laboratory. Using this instrument we profiled thousands of single cells derived from synovial tissue obtained from an RA patient immediately after surgery. This methodology allowed us to profile gene expression in an unbiased and highly quantitative manner across all cell populations simultaneously, providing an attractive alternative to bulk-sorting followed by RNA-seq. Single cell deconvolution of synovial tissue revealed immune subsets including CD4+, CD8+ T cells, and NK cells that likely contribute to RA disease etiology through expression of signaling molecules and their interactions with immune and fibroblast populations. Further, single cell transcriptomic signatures identified sources of heterogeneity, specifically in fibroblasts, corresponding to differences in microenvironment and function. Importantly, this dataset can be used to discover and validate putative markers enabling future functional studies. Here we used immunofluorescence to localize CD55+ and CD90+ synovial fibroblasts to the lining and sublining respectively. More generally this work offers a first comprehensive glimpse into inflamed synovial tissue at the single-cell level and is a step towards compiling an RA cell atlas. Rich datasets such as this could potentially elucidate disease mechanisms in RA and be used to stratify patients, ascertain treatment efficacy, and classify disease sub-states across many arthritic conditions.

## Methods

### Microfluidic chip fabrication

Microfluidic chips were designed in AutoCAD (Autodesk) and a transparency mask was manufactured (Advance Reproductions). SU-8 3050 (MicroChem) was spin coated onto a clean silicon wafer to a thickness of 100 μm and exposed through the mask using a contact mask aligner. After development, polydimethylsiloxane (PDMS) was poured over the master mold, degassed in a desiccator and cured in an oven at 80°C for 2 hours. PDMS slabs were cut from the substrate, holes were punched for tubing connections and the slab was bonded to a clean glass slide using oxygen plasma. Finally, microfluidic channels were treated with Aquapel (Pittsburgh Glass Works) and dried in an oven at 80°C for 30 minutes.

### Microfluidic control instrument

The microfluidic control instrument consists of a 3D printed frame affixed with a custom printed circuit board (PCB) designed in Eagle (Autodesk) containing electronic and pneumatic components. The frame accommodates a microfluidic chip viewed at fixed focus through a Raspberry pi camera with lens. Fluid flow through the microfluidic chip is achieved through pressurization of the head space of reservoir vials situated at the rear of the instrument using a micro air pump, two independent regulators and micro solenoid valves. Pressures for the oil vial and the aqueous vials (cell and microparticle) were independently measured using two analog gauge pressure sensors. Barcoded microparticles were stirred with a stir bar located inside of the vial under the influence of a permanent magnet affixed to a stepper motor shaft situated at the base of the instrument. The instrument is controlled through a Raspberry Pi 2 model B single-board computer with a custom graphical user interface for monitoring of the experiment (through the microscope camera) and control of solenoid valves, micro-air pump, and magnetic stirring. The instrument is powered through an external wall adapter power supply (12V, 3A) through a barrel jack connection mounted on the PCB.

### RA Patient synovial tissue disaggregation

Synovial tissue was collected from an RA patient enrolled and genetically consented under the HSS Early RA Tissue Study (IRB# 2014–317) during a synovectomy procedure. The patient was under 40 years old, seropositive with high titers for CCP antibodies and met 2010 ACR/EULAR Criteria^25^.The HSS Pathology Lab confirmed the sample was synovial tissue by gross inspection and histologic examination of OCT- and Paraffin-embedded blocks. For single-cell suspensions, the synovial tissue was minced with scissors to ~2mm^3^ pieces, which were then digested with Liberase TL (100 μg/mL, Roche) and DNAsel (100ug/mL, Roche) at 37°C for 15 minutes with inversion of the sample every 5 minutes. The enzymatic reaction was quenched by 10% fetal bovine serum in RPMI (Invitrogen) and debris filtered out using two 70 μm strainers. Red blood cells were lysed (reagent a gift of J. Lederer) for 5 min at room temperature, followed by an additional filter step through a 70 μm strainer. The filtration steps should remove large pieces of debris, as well as poorly disaggregated cell clumps. Cells were counted on a hemocytometer and assessed for viability (>85%) using trypan blue staining and 150,000 synoviocytes were re-suspended in Drop-seq loading buffer.

### Single-cell droplet experiments

Single-cell experiments are nearly identical to those described in Macosko *et al*., save for the addition of Ficoll PM-400 to the cell buffer to match fluid viscosity of the aqueous flows. Briefly, cells and split-pool synthesized barcoded microparticles suspended in lysis buffer are co-encapsulated into nanoliter volume droplets. Microparticles contained oligos consisting of a cell barcode (same for all oligos on a microparticle), a UMI (different for each oligo on a microparticle), a PCR handle and a polyT stretch for capture of polyA mRNA. Following encapsulation (and immediate cell lysis) mRNAs hybridize to the microparticle, the emulsion is broken, microparticles are collected and cDNA is generated through reverse transcription in bulk. Exonuclease, PCR (15 cycles total), cDNA purification, and Nextera library preparation steps were performed as in the original manuscript. Libraries were sent for sequencing on the Illumina HiSeq 2500 platform.

### Single-cell RNA-seq analysis

The raw sequencing data were processed as in Macosko *et al.* Briefly, reads were aligned to the UCSC hg19 transcriptome and then binned and collapsed onto the cell barcodes corresponding to individual microparticles using Drop-seq tools (http://mccarrolllab.com/dropseq). To exclude low quality cells, we filtered out cells for which fewer than 500 genes were detected and excluded likely doublets by removing cells with greater than 13,000 UMIs. All genes that were not detected in at least 3 cells were discarded, leaving 24,576 genes. Library-size normalization was performed on the UMI-collapsed gene expression values for each cell barcode by scaling by the total number of transcripts and multiplying by 10,000. The data was then natural-log transformed before any further downstream analysis with Seurat.

For each gene, we constructed a generalized linear regression model (with negative binomial errors) to predict gene expression based on cellular read depth, percentage of mitochondrial genes detected, run ID, and alignment rate to the transcriptome. We used the scaled (z-scored) Pearson residuals from this model as corrected gene expression estimates for downstream dimensional reduction. We first selected 2,117 genes with high variance, using the MeanVarPlot function with log-mean expression values between 0 and 8 and dispersion (variance/mean) between 0.8 and 30. We then reduced the dimensionality of our data using independent component analysis and identified 32 independent components (ICs) for downstream analysis. We removed one IC that was driven primarily by cell cycle genes. We then utilized the smart local moving algorithm for modularity-driven clustering^44^, based on a cell-cell distance matrix constructed on these ICs. This was implemented using the FindClusters function in Seurat with a resolution of 1.6, a k.param of 40, and prune.SNN set to 0.1 to identify 16 distinct clusters of cells.

We and others^5^ have noticed that while modularity-based clustering is a sensitive method for community detection, it can be affected by the multi-resolution problem, and can occasionally over-partition large clusters in order to sensitively detect rare populations. We therefore implemented a post-hoc procedure to merge together clusters with similar gene expression patterns. We reasoned that if a partitioning represented ‘overclustering’ of the data, it would be challenging to distinguish the two resulting cluster based on gene expression values. Therefore, for each pair of clusters, we trained a random forest classifier to predict cluster membership based on the expression level of variable genes, using the ranger package in R with default parameters^45^. We merged clusters together if the classifier had a prediction error greater than 13% as measured by the out-of-bag error. This procedure resulted in the iterative merging of two pairs of clusters, both of which also had few differentially expressed genes between them. For visualization, we applied t-distributed stochastic neighbor embedding (t-SNE) on the cell loadings of the previously selected ICs to view the cells in two dimensions.

Additionally, to explore potential heterogeneity within the NK cell cluster (Supplementary Fig. 2), we took all GNLY+ NK cells and performed a PCA on the 1,000 genes with the highest variance/mean ratio, observing that PC1 and PC2 separated CD56^bright^ from CD45^dim^ NK populations.

### Immunofluorescence

Antibodies for CD55 (NaM16-4D3) and CD90 (EPR3133) were purchased from Santa Cruz Biotechnology, INC. and Abcam, respectively. Sectioning of paraffin-embedded synovial tissue and immunofluorescent staining was performed by the Molecular Cytology Core Facility at Memorial Sloan-Kettering Cancer Center.

### Flow Cytometry

Synovial cell suspensions were stained with fluorochrome-conjugated CD45 (H130Biolegend), CD90 (5E10-Biolegend), CD55 (JS11-Miltenyi Biotec), podoplanin (REA446-Miltenyi Biotec), propidium iodide (PI) (Invitrogen) and analyzed by FACS. Data were analyzed using FlowJo (Tree Star, Inc.) software.

### Identification of differentially expressed genes

To identify marker genes for each cluster, we used the FindAllMarkers command in Seurat, applying the negative binomial differential expression test (‘negbinom’) that we have previously applied to Drop-seq data^46^. Briefly, this test models expression data for individual genes as a generalized linear model with negative binomial errors. The test compares two models with a likelihood ratio test, one constructed with a group (i.e. cluster) indicator variable, and the other assuming an identical model for all clusters. For each cluster, we performed differential expression tests comparing cells within that cluster, to all other cells in the dataset, applying a Bonferroni correction for returned p-values in Supplementary Table 2.

### GO enrichment

We performed pathway and gene-ontology (GO) enrichment for the fibroblast and T cell clusters using the pagoda routines from the scde package on the scaled and normalized scRNA-seq data. We used the genome wide annotation for humans as our reference (Carlson M (2017). *org.Hs.eg.db: Genome wide annotation for Human.* R package version 3.4.1). We performed a PCA analysis and the top principle component for each gene set was obtained using the pagoda.pathway.wPCA function. We then evaluated the statistical significance of each gene set using the pagoda.top.aspects function and retained those with a p-value of less than 0.01. To remove redundant GO terms, we used the pagoda.reduce.loading.redundancy function to collapsed gene sets driven by the same combinations of genes and the pagoda.reduce.redundancy function to collapse those that separated the same sets of cells. Finally, we took the GO terms with the 10 highest average cell PC1 score for each of our identified clusters for heatmap visualization and analysis.

## Acknowledgements

We thank the HSS orthopedic surgeons (particularly Dr. M. Figgie), Dr. Edward DiCarlo of HSS Pathology Department for gross examination and histologic scoring of the synovial sample, rheumatologists, clinical research coordinators (particularly R. Cummings, M. McNamara and S. Mirza), research technicians (particularly J. Ding, I. Cohn), HSS Research Pathologist Dr. Tania Pellegrini, additional staff in the New York Genome Center Technology/Innovation and Satija Labs, and the consenting RA patient who contributed by providing the tissue used in this study. Supported by 5UH2AR067691-03 (VB; RBD; LI, with supplemental funding to RS), and an NIH New Innovator Award (1DP2HG009623-01 to RS).

## Author contributions

WS conceived of, designed, built, and tested the microfluidic control instrument with input from HPS and RS. WS LTD DEO VPB RBD HPS and RS conceived the application to synovial RA tissue. LTD prepared the synovectomy samples and WS AR processed the samples with Drop-seq. AB WS and RS analyzed the single cell RNA-seq data. CR and LTD performed immunofluorescence and FACS experiments. SMG LBI VPB DEO RBD HPS and RS supervised the research, and provided reagents and funding.

---

**Supplementary Figure 1.**
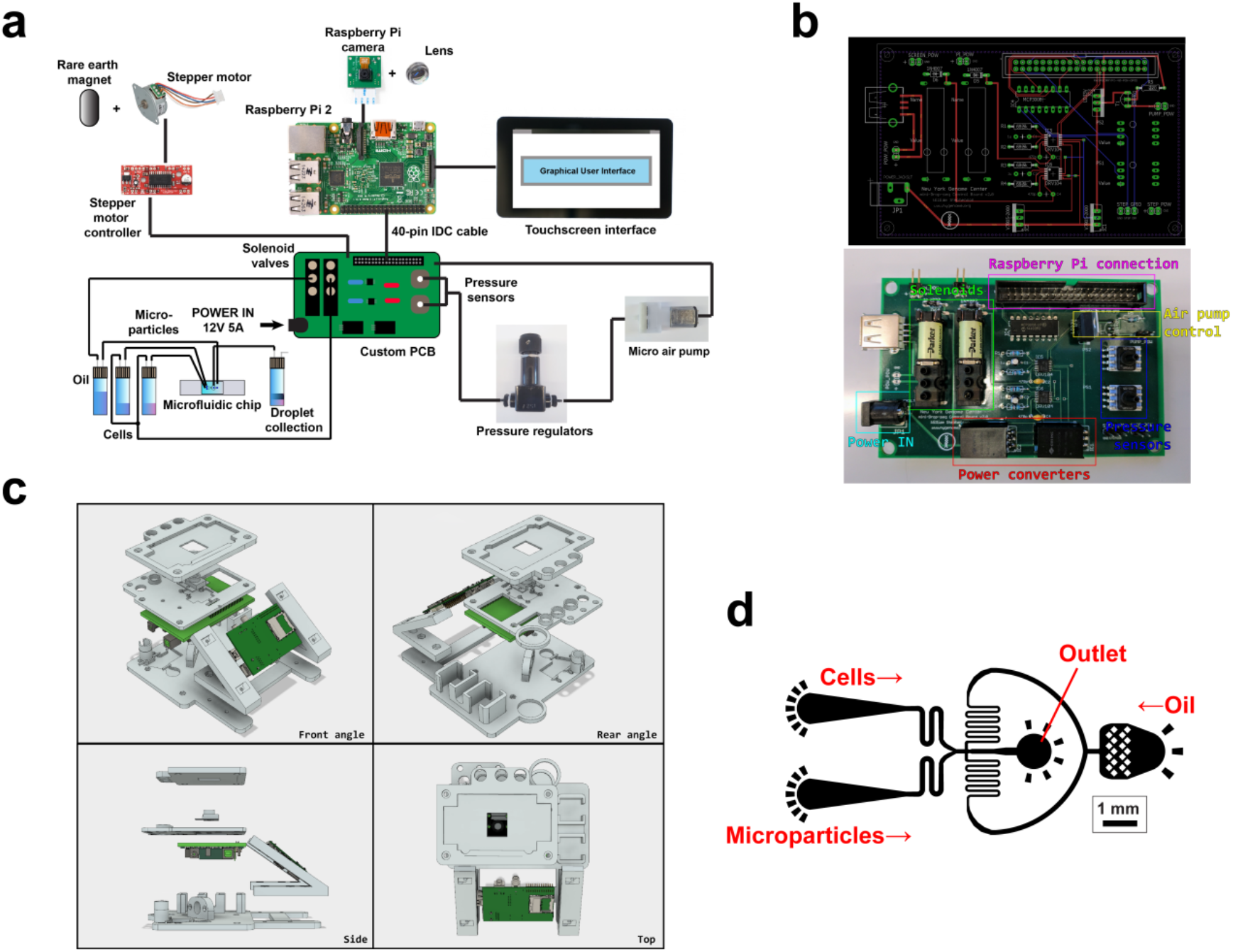
Microfluidic control instrument design and microfluidic chip design. **a**) Component diagram of the instrument. The instrument is controlled with a Raspberry pi 2 model B single board computer that interfaces with the components through a custom designed printed circuit board (PCB). **b**) Circuit layout (top) and image (bottom) of the completed PCB. **c**) Multi-angle view of the 3D printed instrument frame. **d**) Microfluidic chip design. Cell and microparticle inlets have equal hydrodynamic resistance up to the junction with the bifurcated oil channel.

**Supplementary Figure 2.**
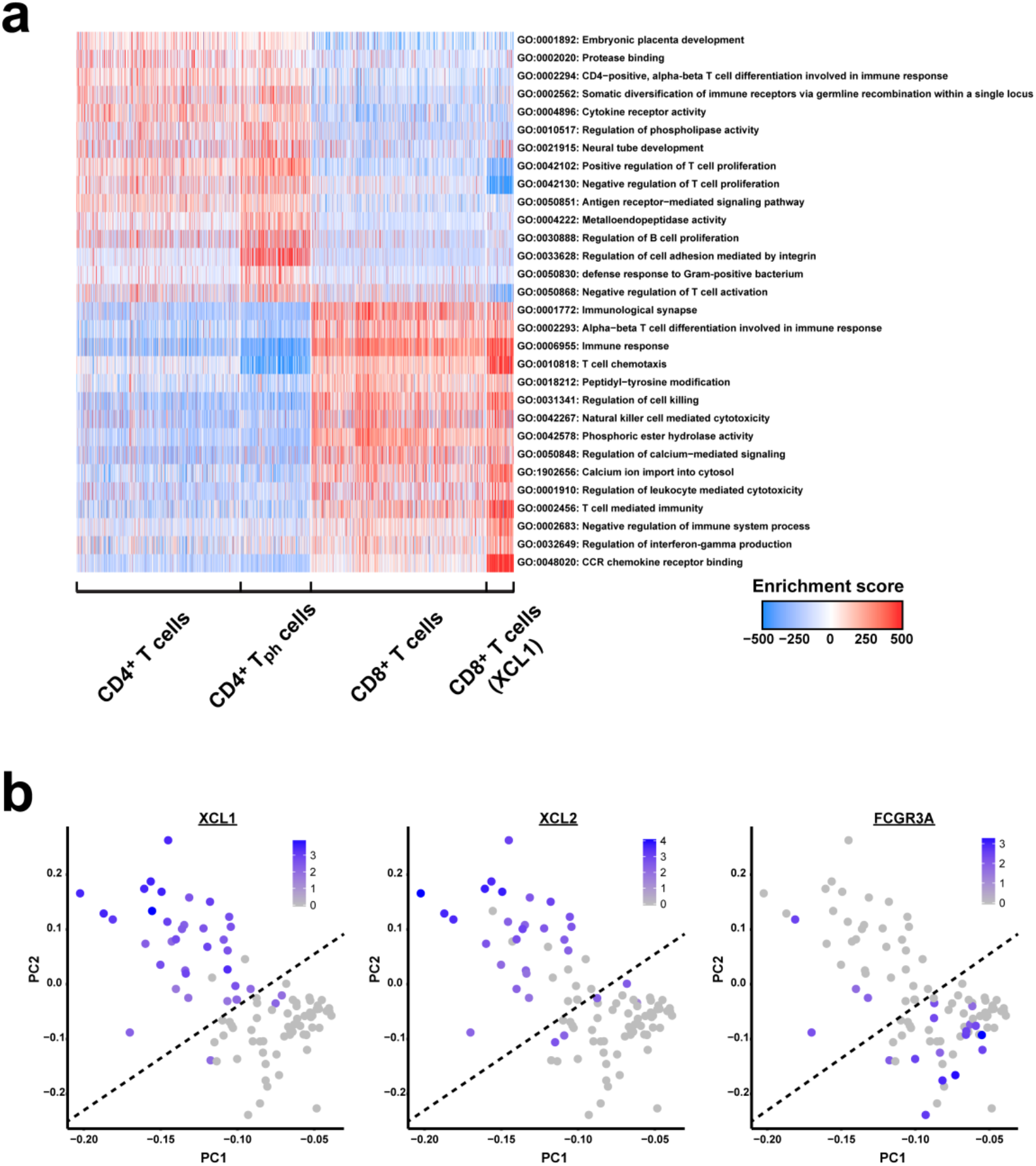
T cell GO enrichment and NK cell subdivision. a) Pathway and gene set overdispersion analysis on the four T cell populations identified via unbiased clustering of the single cell RNA-seq data. The enrichment score corresponds to each cells’ first principle component loading from pathway analysis as computed in *pagoda*. b) Comparison of *XCL1, XCL2*, and *FCGR3A* expression across *GNLY*+ NK cells plotted on the first two principle components after performing a PCA on the top 1,000 genes with the highest variance/mean ratio. The dashed line corresponds to the separation of CD56^bright^ and CD56^dim^ NK cells.

**Supplementary Table 1.**
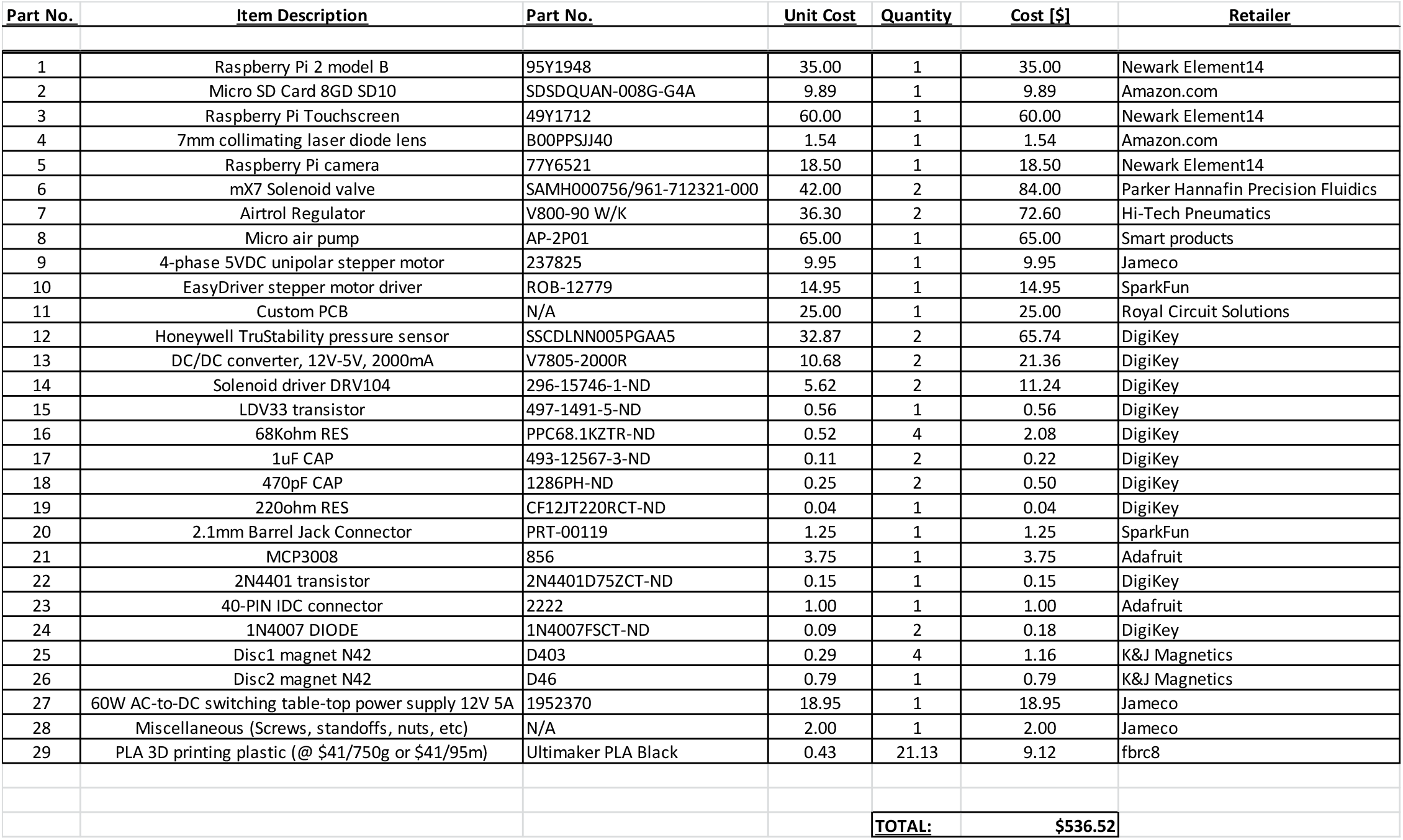
Bill of materials (BOM) for the microfluidic control instrument. Prices as of 5/4/2017.

## References

1. Macosko, E. Z. et al. Highly parallel genome-wide expression profiling of individual cells using nanoliter droplets. Cell 161, 1202–1214 (2015).

2. Klein, A. M. et al. Droplet Barcoding for Single-Cell Transcriptomics Applied to Embryonic Stem Cells. Cell 161, 1187–1201 (2015).

3. Vitak, S. A. et al. Sequencing thousands of single-cell genomes with combinatorial indexing. Nat. Methods 14, (2017).

4. Zheng, G. X. Y. et al. Massively parallel digital transcriptional profiling of single cells. Nat. Commun. 8, 1–12 (2017).

5. Shekhar, K. et al. Comprehensive Classification of Retinal Bipolar Neurons by Single-Cell Transcriptomics. Cell 166, 1308–1323.e30 (2016).

6. Baron, M. et al. A Single-Cell Transcriptomic Map of the Human and Mouse Pancreas Reveals Inter- and Intra-cell Population Structure. Cell Syst. 3, 346–360 (2016).

7. Shembekar, N., Chaipan, C., Utharala, R. & Merten, C. A. Droplet-based microfluidics in drug discovery, transcriptomics and high-throughput molecular genetics. Lab Chip (2016). doi:10.1039/C6LC00249H

8. Kaminski, T. S., Scheler, O. & Garstecki, P. Droplet microfluidics for microbiology: techniques, applications and challenges. Lab Chip 16, 2168–2187 (2016).

9. Alles, J. et al. Cell fixation and preservation for droplet-based single-cell transcriptomics. bioRxiv (2017).

10. Gierahn, T. M. et al. Seq-Well: portable, low-cost RNA sequencing of single cells at high throughput. Nat. Methods 1–8 (2017). doi:10.1038/nmeth.4179

11. Yuan, J. & Sims, P. A. An Automated Microwell Platform for Large-Scale Single Cell RNA-Seq. bioRxiv 70193 (2016). doi:10.1101/070193

12. Bose, S. et al. Scalable microfluidics for single-cell RNA printing and sequencing. Genome Biol. 16, 120 (2015).

13. Briggs, A. W. et al. Tumor-infiltrating immune repertoires captured by single-cell barcoding in emulsion. bioRxiv 1–34 (2017). at <http://biorxiv.org/content/early/2017/05/05/134841.abstract>

14. Stoeckius, M. et al. Large-scale simultaneous measurement of epitopes and transcriptomes in single cells. bioRxiv (2017). doi:10.1101/113068

15. Zhang, Y. & Jiang, H. R. A review on continuous-flow microfluidic PCR in droplets: Advances, challenges and future. Anal. Chim. Acta 914, 7–16 (2016).

16. Sidore, A. M., Lan, F., Lim, S. W. & Abate, A. R. Enhanced sequencing coverage with digital droplet multiple displacement amplification. Nucleic Acids Res. gkv1493 (2015). doi:10.1093/nar/gkv1493

17. Kamperman, T., Henke, S., Visser, C. W., Karperien, M. & Leijten, J. Centering Single Cells in Microgels via Delayed Crosslinking Supports Long-Term 3D Culture by Preventing Cell Escape. Small 1603711 (2017). doi:10.1002/smll.201603711

18. Siltanen, C. et al. One Step Fabrication of Hydrogel Microcapsules with Hollow Core for Assembly and Cultivation of Hepatocyte Spheroids. Acta Biomater. 50, 428–436 (2017).

19. Lee, K. et al. Generalized serial dilution module for monotonic and arbitrary microfluidic gradient generators. Lab Chip 9, 709–17 (2009).

20. Ozkumur, E. et al. Inertial focusing for tumor antigen-dependent and -independent sorting of rare circulating tumor cells. Sci. Transl. Med. 5, 179ra47 (2013).

21. Martel, J. M. et al. Continuous Flow Microfluidic Bioparticle Concentrator. Sci. Rep. 5, 11300 (2015).

22. Di Carlo, D., Irimia, D., Tompkins, R. G. & Toner, M. Continuous inertial focusing, ordering, and separation of particles in microchannels. Proc. Natl. Acad. Sci. U. S. A. 104, 18892–18897 (2007).

23. Scott, D. L., Wolfe, F. & Huizinga, T. W. J. Rheumatoid arthritis. Lancet 376, 1094–1108 (2010).

24. McInnes, I. B. & Schett, G. The Pathogenesis of Rheumatoid Arthritis. N. Engl. J. Med. 365, 2205–19 (2011).

25. Aletaha, D. et al. 2010 Rheumatoid arthritis classification criteria: An American College of Rheumatology/European League Against Rheumatism collaborative initiative. Arthritis Rheum. 62, 2569–2581 (2010).

26. Buckley, C. D., Barone, F., Nayar, S., Bénézech, C. & Caamaño, J. Stromal cells in chronic inflammation and tertiary lymphoid organ formation. Annu. Rev. Immunol. 33, 715–45 (2015).

27. Noss, E. H. & Brenner, M. B. The role and therapeutic implications of fibroblast-like synoviocytes in inflammation and cartilage erosion in rheumatoid arthritis. Immunol. Rev. 223, 252–270 (2008).

28. Naylor, A. J., Filer, A. & Buckley, C. D. The role of stromal cells in the persistence of chronic inflammation. Clin. Exp. Immunol. 171, 30–35 (2013).

29. Bartok, B. & Firestein, G. S. Fibroblast-like synoviocytes: key effector cells in rheumatoid arthritis. 233, 233–255 (2010).

30. Fassbender, H. G. Histomorphological basis of articular cartilage destruction in rheumatoid arthritis. Coll. Relat. Res. 3, 141–155 (1983).

31. Müller-Ladner, U. et al. Synovial fibroblasts of patients with rheumatoid arthritis attach to and invade normal human cartilage when engrafted into SCID mice. Am. J. Pathol. 149, 1607–15 (1996).

32. Lafyatis, R. et al. Anchorage-independent Growth of Synoviocytes from Arthritic and Normal Joints. J. Clin. Investig. Inc. 83, 1267–1276 (1989).

33. Müller-ladner, U., Ospelt, C., Gay, S., Distler, O. & Pap, T. Cells of the synovium in rheumatoid arthritis Synovial fibroblasts. 10, 1–10 (2007).

34. Pap, T. et al. Activation of synovial fibroblasts in rheumatoid arthritis: lack of Expression of the tumour suppressor PTEN at sites of invasive growth and destruction. Arthritis Res. 2, 59–64 (2000).

35. Pap, T., Müller-Ladner, U., Gay, R. E. & Gay, S. Fibroblast biology. Role of synovial fibroblasts in the pathogenesis of rheumatoid arthritis. Arthritis Res. 2, 361–7 (2000).

36. Gravallese, E. M., Darling, J. M., Ladd, A. L., Katz, J. N. & Glimcher, L. H. In situ hybridization studies of stromelysin and collagenase messenger RNA expression in rheumatoid synovium. Arthritis Rheum. 34, 1076–1084 (1991).

37. Villani, A.-C. et al. Single-cell RNA-seq reveals new types of human blood dendritic cells, monocytes, and progenitors. Science (80-.). 356, eaah4573 (2017).

38. Proof, G. et al. Pathologically expanded peripheral T helper cell subset drives B cells in rheumatoid arthritis. Nat. Publ. Gr. 542, 110–114 (2017).

39. Fan, J. et al. Characterizing transcriptional heterogeneity through pathway and gene set overdispersion analysis. Nat. Methods 13, 241–244 (2016).

40. Blaschke, S. et al. Expression of activation-induced, T cell-derived, and chemokine-related cytokine/lymphotactin and its functional role in rheumatoid arthritis. Arthritis Rheum. 48, 1858–1872 (2003).

41. Borthwick, N. J. et al. Selective migration of highly differentiated primed T cells, defined by low expression of CD45RB, across human umbilical vein endothelial cells: effects of viral infection on transmigration. Immunology 90, 272–80 (1997).

42. Poli, A. et al. CD56bright natural killer (NK) cells: An important NK cell subset. Immunology 126, 458–465 (2009).

43. Jurisic, G., Iolyeva, M., Proulx, S. T., Halin, C. & Detmar, M. Thymus cell antigen1 (Thy1, CD90) is expressed by lymphatic vessels and mediates cell adhesion to lymphatic endothelium. Exp. Cell Res 316, (2010).

44. Waltman, L. & Van Eck, N. J. A smart local moving algorithm for large-scale modularity-based community detection. Eur. Phys. J. B 86, 1–33 (2013).

45. Wright, M. N. & Ziegler, A. ranger: A Fast Implementation of Random Forests for High Dimensional Data in C++ and R. arXiv.org stat.ML, (2015).

46. Mayer, C. et al. Inhibitory neuron diversity originates from cardinal classes shared across germinal zones. Bioarxiv 1–3 (2017).

